# Effector-dependent activation and oligomerization of NRC helper NLRs by Rpi-amr3 and Rpi-amr1

**DOI:** 10.1101/2022.04.25.489359

**Authors:** Hee-Kyung Ahn, Xiao Lin, Andrea Carolina Olave-Achury, Lida Derevnina, Mauricio P Contreras, Jiorgos Kourelis, Sophien Kamoun, Jonathan D G Jones

**Author notes:** These authors contributed equally (Hee-Kyung Ahn, Xiao Lin).

## Abstract

Plant pathogens compromise crop yields. Plants have evolved robust innate immunity that depends in part on intracellular Nucleotide-binding, Leucine Rich-Repeat (NLR) immune receptors that activate defense responses upon detection of pathogen-derived effectors. Most “sensor” NLRs that detect effectors require the activity of “helper” NLRs, but how helper NLRs support sensor NLR function is poorly understood. Many Solanaceae NLRs require the NRC (NLR-Required for Cell death) class of helper NLRs. We show here that Rpi-amr3, a sensor NLR from *Solanum americanum*, detects AVRamr3 from the potato late blight pathogen, *Phytophthora infestans*, and activates oligomerization of the helper NLR NRC2 into a high-molecular weight resistosome. The NRC2 resistosome also forms upon recognition of *P. infestans* effector AVRamr1 by another sensor NLR, Rpi-amr1. The ATP-binding motif of Rpi-amr3 is required for NRC2 resistosome formation, but not for interaction with the cognate effector. The NRC2 resistosome can be activated by AVRamr3 homologs from other *Phytophthora* species. Mechanistic understanding of NRC resistosome formation will underpin engineering crops with durable disease resistance.

## Introduction

Plants have powerful defense mechanisms, but to be effective, these must be rapidly activated at sites of attempted pathogen ingress. Activation of defense requires detection, both by cell surface receptors that usually detect pathogen-derived components such as flagellin or chitin (Lee *et al*, 2021), and by intracellular Nucleotide-binding, Leucine-rich Repeat (NLR) receptors which detect effectors that often function for the pathogen to attenuate plant defenses (Jones *et al*, 2016).

Natural plant populations carry extensive genetic variation in immune receptor repertoires (Ngou *et al*, 2022b). Plant breeders have long exploited this genetic variation to elevate crop varietal resistance by introgression of multiple disease *Resistance* (*R*) genes from wild relatives. *R* genes usually encode NLR immune receptors (Meyers *et al*, 1999). Some plant species carry scores or even hundreds of different NLR immune receptor genes, with extensive allelic diversity and presence/absence polymorphism (Barragan & Weigel, 2021). These NLR immune receptors can confer resistance to bacteria, fungi, oomycetes, viruses and even invertebrates (Ngou *et al*, 2022a), triggering broad interest in how these immune receptors can activate defense upon recognition of molecules from such diverse sources.

NLRs are broadly categorized into three subclasses, based on their N-terminal domains. TIR-NLRs, CC-NLRs, and CC_R_-NLRs have N-terminal Toll-like, Interleukin-1 receptor, Resistance (TIR) domains, Coiled-Coil (CC) domains, and RPW8-like (CC_R_) domains, respectively (Collier *et al*, 2011; Meyers *et al*., 1999). The N-terminal domains of the NLRs have direct roles in signaling upon effector-dependent oligomerization of the NLRs. For example, the TIR-NLRs ROQ1 and RPP1 form a tetramer upon detection of their cognate recognized effectors, activating an NADase activity by forming a tetramer of the TIR domain (Ma *et al*, 2020; Martin *et al*, 2020). The enzymatic activity generates small molecules that are required for downstream signaling via EDS1 (Huang *et al*, 2022). The oligomerization of ZAR1 and Sr35 into a pentamer upon effector detection induces assembly of the α-helices in the CC domains into a cation channel in the plasma membrane that is required for signalling (Bi *et al*, 2021; Förderer *et al*,2022; Wang *et al*, 2019a). The CC_R_-NLR NRG1.1 and ADR1 also oligomerize and localize to the plasma membrane and can mediate calcium ion influx (Jacob *et al*, 2021). However, whether all NLRs form resistosomes upon activation is unclear.

NLRs often function independently as singletons, but accumulating evidence suggests that many NLRs function in pairs or networks. The effector-detecting NLR is often named a “sensor” NLR, whereas the downstream signaling NLRs that convert recognition into immune activation are called “helper” NLRs (Feehan *et al*, 2020). Paired NLRs, such as RRS1/RPS4 or RGA4/RGA5, are divergently transcribed, and one of the NLRs often carries an integrated domain (ID) for effector detection, while the other signals upon recognition (Cesari *et al*, 2014). In contrast, helper NLRs of the CC_R_-NLR and NRC classes usually map to different genomic loci from the “sensor” NLRs and are required for the activity of multiple “sensor” NLRs (Feehan *et al*., 2020; Jubic *et al*, 2019; Wu *et al*, 2018). The first CCR-type helper NLR, NRG1 (Peart *et al*, 2005), was found to be required for the function of TIR-NLRs (Castel *et al*, 2019; Qi *et al*,2018; Wu *et al*, 2019), and the related ADR1 helper NLRs can contribute to both TIR-NLR and CC-NLR function (Saile *et al*, 2020).

The NRC class of helper NLR was discovered in the Solanaceae and is widespread in the asterid but not the rosid clade of angiosperms (Wu *et al*, 2017). The NRC class of helper NLRs are phylogenetically related to their corresponding sensor NLRs in the asterid plant family, and ~50% of Solanaceae NLRs are either NRC-dependent sensor NLRs or NRCs (Wu *et al*., 2017). Different sensor NLRs depend on different combinations of helper NLRs to activate the immune response (Wu *et al*., 2018). A conserved N-terminal MADA motif was found in the NRC family and in ~20% of other CC-NLRs, including ZAR1 from Arabidopsis (Adachi *et al*, 2019), leading to the hypothesis that NRCs might activate defense via similar mechanisms to ZAR1. Mutating the MADA motifs of NRC2, NRC3 or NRC4 from *Nicotiana benthamiana* results in loss of function, and the corresponding N-terminal α1 helix can be swapped with the equivalent region of ZAR1 (Adachi *et al*., 2019; Duggan *et al*, 2021; Kourelis *et al*, 2021). This suggests that the MADA motif of NRCs might form a cation-selective channel to activate immune signalling and cell death, as does ZAR1 (Bi *et al*., 2021). However, how the NRC-dependent immune response is activated upon effector detection by sensor NLRs remains unknown.

*Phytophthora* diseases cause yield loss for many important crop plants (Kamoun *et al*, 2015). These diseases are mainly controlled by agrichemical sprays (Cooke *et al*, 2011). Many *R* genes against *P. infestans* (*Rpi*) genes were cloned from wild potatoes (Vleeshouwers *et al*,2011). We cloned the *Rpi-amr3* and *Rpi-amr1* genes from *Solanum americanum* for resistance against *P. infestans*, and both *Rpi* genes confer broad-spectrum resistance against late blight in potato (Lin *et al*, 2021; Witek *et al*, 2016; Witek *et al*, 2021). We also defined the cognate effectors *Avramr3* and *Avramr1* from *P. infestans* (Lin *et al*., 2021; Lin *et al*, 2020). Rpi-amr3 and Rpi-amr1 can also recognize AVRamr3 and AVRamr1 homologs from other *Phytophthora* pathogens (Lin *et al*., 2021; Witek *et al*., 2021). Rpi-amr3 and Rpi-amr1 are NRC2/3/4- and NRC2/3-dependent, respectively (Lin *et al*., 2021; Witek *et al*., 2021).

Here, we used *nrc2/3/4* CRISPR knockout (*KO*) *N. benthamiana* line (Wu *et al*, 2020) and transient expression of Rpi-amr3/AVRamr3 or Rpi-amr1/AVRamr1, and of a NRC2 mutant in its MADA motif (NRC2^EEE^) to study the activation of the helper NLR NRC2. We found that upon effector recognition, both Rpi-amr3 and Rpi-amr1 activate formation of a high-molecular weight complex of NRC2, dependent on a functional ATP-binding motif in the Rpi-amr protein. Intriguingly, some AVRamr3 homologs from other *Phytophthora* pathogens such as *P. parasitica* can also activate NRC2 resistosome formation through Rpi-amr3. This finding could be pivotal for breaking the restricted taxonomic functionality (RTF) of some *NLR* genes and elevating disease resistance in crops that lack *NRC* genes (Tai *et al*, 1999).

## Results

### Rpi-amr3 and AVRamr3 form an NRC-independent protein complex

Rpi-amr3 and Rpi-amr1 are canonical CC-NLRs in *S. americanum* that recognize *P. infestans* effectors AVRamr3 and AVRamr1, respectively (Lin *et al*., 2021; Lin *et al*., 2020; Witek *et al*., 2016; Witek *et al*., 2021) (Fig 1A). Both AVRamr3 (Lin *et al*., 2021) and AVRamr1 are RXLR effectors with predicted WY domains (Fig EV1A-D) (Boutemy *et al*, 2011). Multiple NRC NLRs in *N. benthamiana* (*Nb*) support immune activation by Rpi-amr3 and Rpi-amr1 (Lin *et al*., 2021; Witek *et al*., 2021) (Fig EV2A). Therefore, we chose transient *Agrobacterium* infiltration in *N. benthamiana* as a model system to test the immune activation mechanism of Rpi-amr3 or Rpi-amr1 via NRC2.

**Figure 1.**
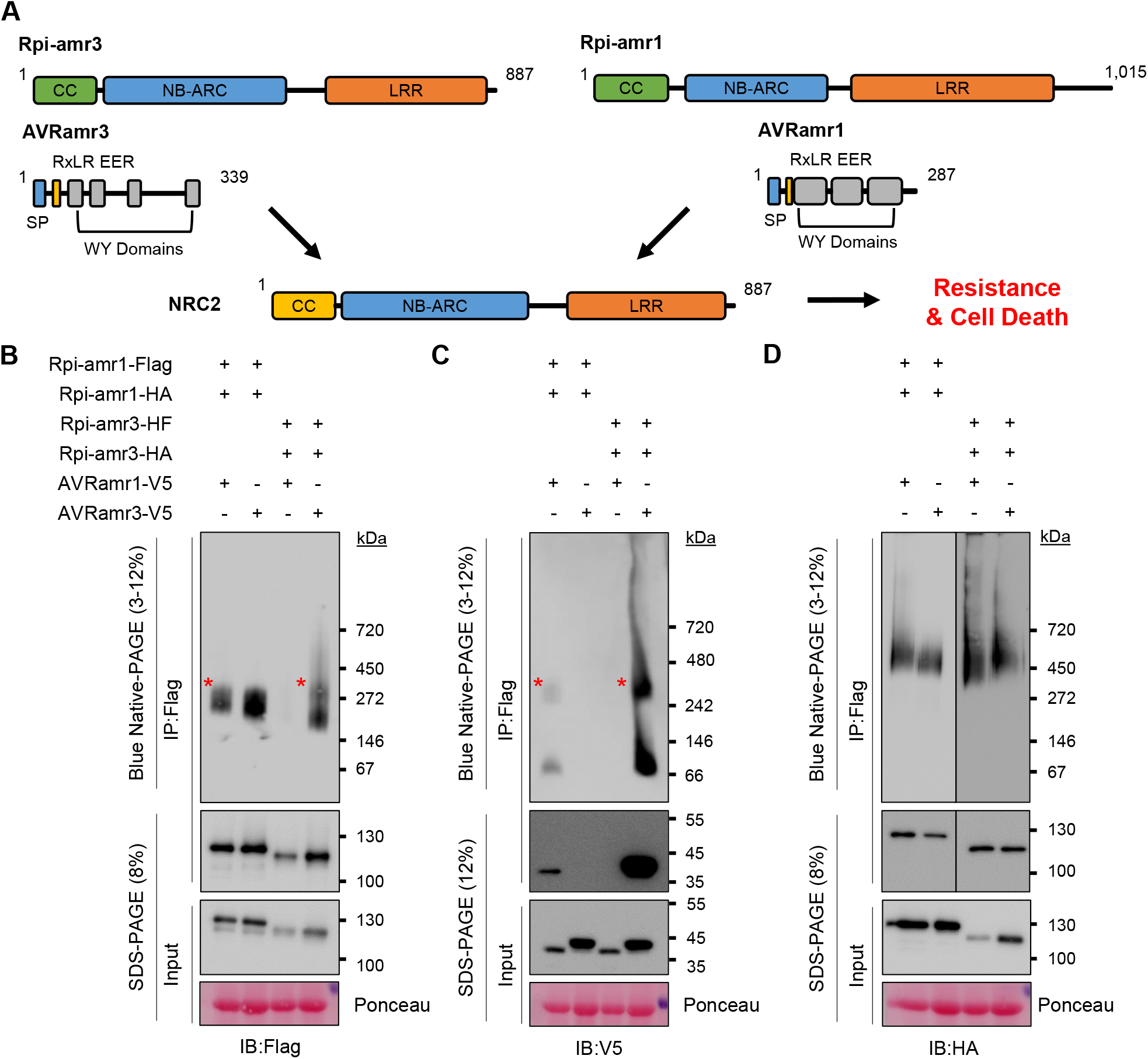
Rpi-amr1 and Rpi-amr3 form a protein complex with AVRamr1 and AVRamr3, respectively. A. Schematic model of NRC-dependent resistance by sensor NLRs Rpi-amr1, Rpi-amr3 and cognate effectors AVRamr1 and AVRamr3, respectively. Each domain is labelled and represented with a different color. CC, coiled-coil; NB-ARC, nucleotide binding domain shared by APAF-1, R genes, CED-4; LRR, Leucine-rich repeat; SP, signal peptide. B. Rpi-amr1 and Rpi-amr3 form protein complex(es) *in vivo*. Protein extracts from *N. benthamiana nrc2/3/4 KO* plants were immunoprecipitated with anti-Flag antibodies and loaded on blue native-PAGE. Rpi-amr3 co-migrating with AVRamr3 is indicated (*). C. AVRamr1 and AVRamr3 form a protein complex with Rpi-amr1 and Rpi-amr3, respectively. Anti-FLAG immunoprecipitated samples from Fig 1B were detected for AVRamr1-V5 and AVRamr3-V5. AVRamr1 co-migrating with Rpi-amr1 and AVRamr3 co-migrating with Rpi-amr3 is indicated (*). D. Rpi-amr1 and Rpi-amr3 can self-associate *in vivo*. Flag-tag immunoprecipitated samples from Fig 1B were visualized for Rpi-amr1-HA and Rpi-amr3-HA. Data information: SDS-boiled input protein extract and IP eluate samples were loaded onto SDS-PAGE as control. Ponceau staining serve as loading control. Molecular weight markers are shown on the right. Experiments were done at least three times with similar results.

Previously, we reported that Rpi-amr3 and AVRamr3 associate *in planta* (Lin *et al*, 2021), but whether Rpi-amr1 associates with AVRamr1 was unknown. Additionally, how Rpi-amr3 and Rpi-amr1 activate immunity upon effector recognition is also unknown. First, we tested whether Rpi-amr3/AVRamr3 and Rpi-amr1/AVRamr1 associate to form high-molecular weight protein complexes without NRC. Flag/HisFlag (HF)- and HA-tagged Rpi-amr1/3 was co-expressed with V5-tagged AVRamr1/3 respectively (Fig 1B-D) in *nrc2/3/4* knockout *N. benthamiana* leaves to avoid a hypersensitive response (HR) that would impair biochemical analysis (Wu *et al*., 2020). Then, immunoprecipitates of Rpi-amr1-Flag and Rpi-amr3-HF were analyzed with blue native-PAGE to visualize changes in protein complex formation.

We used non-denaturing PAGE methods to monitor the presence of protein complexes that differ in size compared to monomers. The use of Coomassie G250 dye in blue native-PAGE enables protein complexes to migrate towards the anode according to their size (Wittig *et al*, 2010). This method was used to identify composition of mitochondrial membrane protein complexes and photosynthetic protein complexes in plants (Eubel *et al*, 2005). Recently, this method has been widely used to identify oligomeric changes of NLRs in plants (Hu *et al*, 2020; Jacob *et al*., 2021; Li *et al*, 2019; Na Ayutthaya *et al*, 2020). We used the blue native-PAGE method to investigate not only protein-protein interactions, but also the approximate size of protein complexes *in vivo*. Rpi-amr1-Flag migrated to one major form independent of the cognate effector AVRamr1 (Fig 1B). Rpi-amr1-Flag co-immunoprecipitated AVRamr1-V5, but not AVRamr3-V5 (Fig 1C). We also tested interaction between Rpi-amr1 and AVRamr1 using split-luciferase assays, which showed that Rpi-amr1 specifically interacts with AVRamr1, but not with AVRamr3 (Fig EV2A). When Rpi-amr1-immunoprecipitated eluates were loaded onto blue native-PAGE, two species of AVRamr1 could be detected. As AVRamr1 interacts with Rpi-amr1, the slow-migrating form of AVRamr1 is likely in a protein complex with Rpi-amr1. However, dissociation may occur after immunoprecipitation, which would explain an AVRamr1 signal at ~60 kDa (Fig 1C).

Rpi-amr3-HF expression was stabilized upon co-expression with cognate effector AVRamr3 (Fig 1B). We detected a slow-migrating protein form of Rpi-amr3-HF when AVRamr3-V5 was co-expressed (Fig 1B, Fig EV3, red asterisk). AVRamr3-V5 that had been co-immunoprecipitated with Rpi-amr3-HF migrated as two different species on blue native-PAGE (Fig 1C, Fig EV3B). However, when AVRamr3 was expressed alone, it migrated on the blue native-PAGE predominantly at ~60 kDa (Fig EV4). This shows that AVRamr3 forms a protein complex with Rpi-amr3 that migrates slower than either Rpi-amr3 and AVRamr3 alone. However, neither slow-migrating forms were of the size expected of an Rpi-amr pentamer in complex with a cognate effector, in contrast to the size of the ZAR1 resistosome (Hu *et al*., 2020).

As Rpi-amr1 and Rpi-amr3 migrate slower than the monomer molecular weight of ~120 kDa in the absence of cognate effector, we tested if Rpi-amr1 and Rpi-amr3 self-associate. We co-expressed Rpi-amr1 or Rpi-amr3 fused with two different tags, Flag or HA, immunoprecipitated Rpi-amr1-Flag or Rpi-amr3-HF and detected HA-tagged Rpi-amr signal (Fig 1D). Our results revealed that both Rpi-amr1 and Rpi-amr3 have the capacity to self-associate. However, in comparison to the majority of Rpi-amr1-Flag and Rpi-amr3-Flag protein that migrates with a size of ~270 kDa, the co-immunoprecipitated Rpi-amr1-HA or Rpi-amr3-HA migrates slower in blue native-PAGE at a size above ~450kDa (Fig 1B, D). This indicates that the majority of the Rpi-amr1 or Rpi-amr3 protein does not self-associate *in vivo*. Furthermore, self-associating Rpi-amr1 and Rpi-amr3 migrate with larger mass than the Rpi-amr1/AVRamr1 or Rpi-amr3/AVRamr3 complexes (Fig 1C, D). Thus, the presence or absence of effector did not change self-association of Rpi-amr1 and Rpi-amr3 (Fig 1D). This is consistent with effector-independent self-association observed for other CC-NLRs, such as RPM1 and MLA1 (El Kasmi *et al*, 2017; Maekawa *et al*, 2011). Therefore, we conclude that the majority of Rpi-amr1 and Rpi-amr3 exist as monomers that form heterodimers with AVRamr1 or AVRamr3. However, additional interactors with these proteins cannot be excluded. The role (if any) of self-associated Rpi-amr1 or Rpi-amr3 remains unclear.

### NRC2 oligomerizes upon AVRamr3-dependent activation of Rpi-amr3 and AVRamr1-dependent activation of Rpi-amr1

Next, we tested whether NRCs oligomerize upon Rpi-amr3 and Rpi-amr1 activation. NRC2 and NRC3 from *N. benthamiana* can both support Rpi-amr3 and Rpi-amr1, but NRC2 proteins express better in transient assays compared to NRC3 (Derevnina *et al*, 2021). Therefore, we used NRC2 from *N. benthamiana*. However, co-expression of NRC2-Myc with Rpi-amr3-HF and AVRamr3-V5 in the *nrc2/3/4 N. benthamiana KO* mutant led to HR in plants and subsequent degradation of Rpi-amr3 and AVRamr3, which precluded further analysis (Fig EV3). To abolish the cell death, we generated an NRC2 MADA mutant, in which the conserved leucine residues (L9, L13, L17) of NRC2 were mutated into glutamates (NRC2^EEE^); this NRC2^EEE^ mutant does not lead to HR or protein degradation when co-expressed with Rpi-amr3/AVRamr3 in *nrc2/3/4 KO N. benthamiana* (Fig EV3). This enabled us to use NRC2^EEE^-Myc to study the biochemical changes resulting from AVRamr3-dependent Rpi-amr3 activation. Interestingly, co-expression of NRC2^EEE^-Myc did not alter protein migration patterns of Rpi-amr3-HF or AVRamr3-V5 on blue native-PAGE (Fig 2A, Fig EV3B). The interaction between Rpi-amr3-HF and AVRamr3-V5 was also independent of NRC2^EEE^-Myc co-expression (Fig 2B, Fig EV3A). Next, using protein lysates of Fig 2A, we observed that, in the pre-activation state, NRC2 migrates as a single protein species. However, upon co-expression with Rpi-amr3 and AVRamr3, there was a pronounced shift in migration of NRC2 protein to slower migrating form of ~ 900 kDa (Figure 2C, red asterisk). There were no changes in the size of NRC2^EEE^-Myc protein itself in SDS-PAGE gels (Fig 2C). This suggests that NRC2 proteins undergo protein oligomerization when Rpi-amr3 recognize AVRamr3.

**Figure 2.**
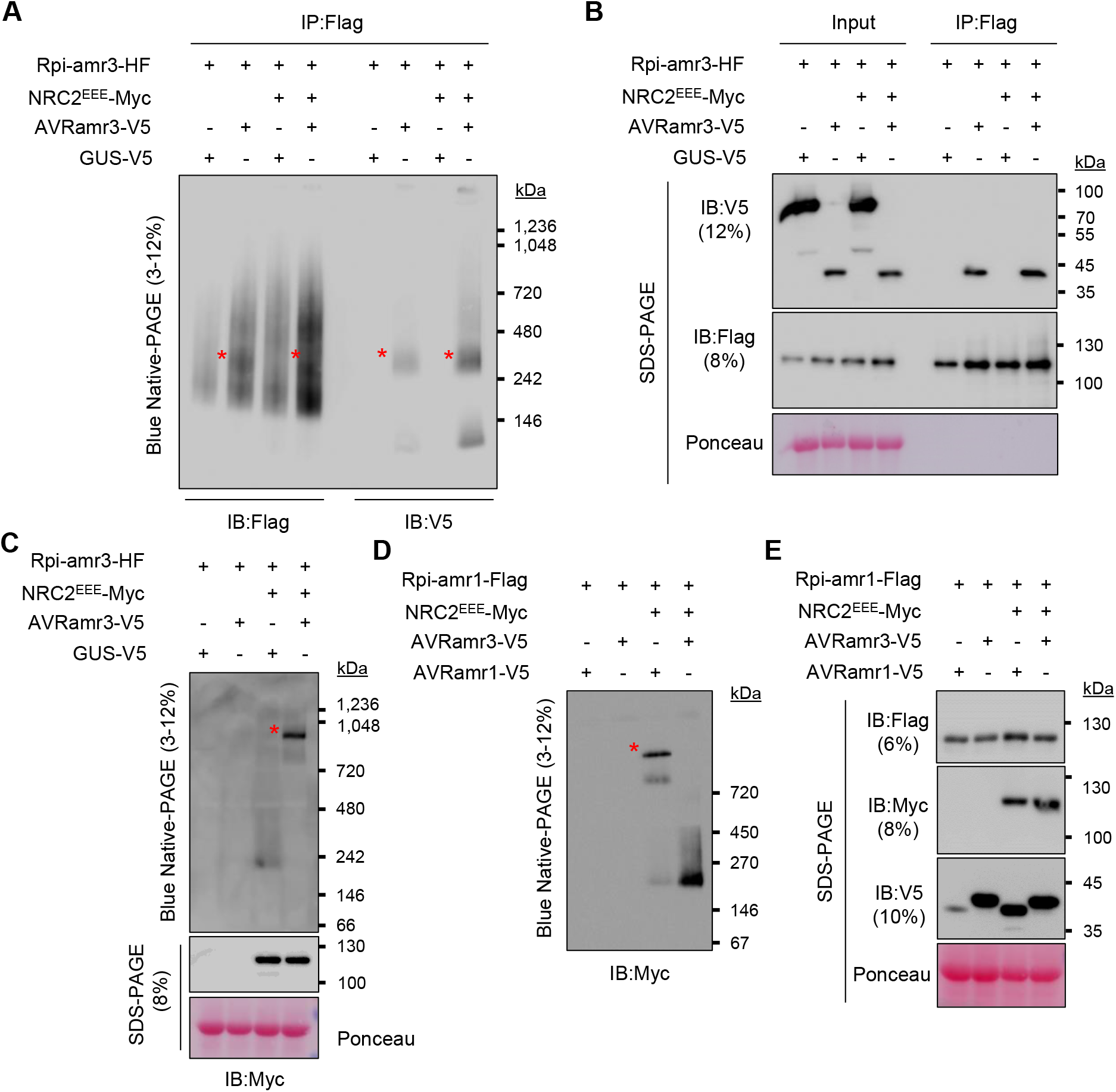
NRC2^EEE^ oligomerizes upon effector detection by Rpi-amr3 and Rpi-amr1. A. NRC2^EEE^-Myc does not change protein complex formation of Rpi-amr3 and AVRamr3. Blue native-PAGE loading of protein extracts from *nrc2/3/4 KO N. benthamiana* plants after immunoprecipitation with anti-Flag antibody. Co-migration of Rpi-amr3-HF and AVRamr3-V5 are indicated (*). Same samples were loaded twice on a blue native-PAGE gel, transferred into one membrane, and only the immunoblotting step was performed separately. B. NRC2^EEE^-Myc does not alter association between Rpi-amr3 and AVRamr3. Samples of Fig 2A were SDS-boiled and loaded on SDS-PAGE. C. NRC2^EEE^-Myc is oligomerized upon effector-dependent activation of Rpi-amr3. Protein lysates from Fig 2A were loaded on blue native-PAGE. SDS-boiled protein lysate samples serve as control for actual size of NRC2^EEE^-Myc. Oligomerized NRC2^EEE^-Myc is indicated (*). D. NRC2^EEE^-Myc oligomerizes upon effector-dependent activation of Rpi-amr1. Protein lysates from *nrc2/3/4* knockout *N. benthamiana* plants were loaded on blue native-PAGE. Oligomerized NRC2^EEE^-Myc is indicated (*). E. Samples from Fig 2D were SDS-boiled and loaded on SDS-PAGE. Protein accumulation of Rpi-amr1-Flag, NRC2^EEE^-Myc, AVRamr1-V5 and AVRamr3-V5 are shown. Data information: Ponceau staining serve as loading control for panels B, C, and E. Molecular weight markers are shown on the right. Experiments were done at least three times with similar results.

To determine whether this phenomenon occurs with other sensor NLRs, we tested migration change of NRC2 in blue native-PAGE upon Rpi-amr1 activation. As with Rpi-amr3/AVRamr3-mediated NRC2 oligomerization, we observed high molecular weight complexes of NRC2^EEE^-Myc in the presence of AVRamr1 and Rpi-amr1, but not in the presence of AVRamr3 and Rpi-amr1, a non-cognate effector of Rpi-amr1 (Fig 2D, E). Thus, we conclude that although Rpi-amr3 and Rpi-amr1 are different sensor NLRs that recognize different effectors, AVRamr3 and AVRamr1, respectively, the resulting change of NRC2 is strikingly similar. In the companion paper by Contreras *et al*., distinct sensor NLRs such as Bs2 and Rx also induce similar changes in NRC2 protein migration on blue native-PAGE.

To confirm that the high-molecular weight complex of NRC2 observed upon Rpi-amr1/AVRamr1 or Rpi-amr3/AVRamr3-mediated activation was indeed NRC2, we conducted 2-dimensional PAGE. We analysed protein extracts from Agro-infiltrated *N. benthamiana* leaf samples on blue native-PAGE as a first dimension, and then performed SDS-PAGE as a second dimension analysis to dissociate protein complexes into individual protein components (Fig 3A). Without Rpi-amr3 activation, most of the NRC2^EEE^ protein migrated faster to ~240 kDa, and migration patterns were similar for Rpi-amr3, with the majority of the NRC2^EEE^ and Rpi-amr3 proteins detected in the faster migrating portion (Fig 3B). When AVRamr3 activated Rpi-amr3, NRC2^EEE^-Myc migrated with a significant shift to ~900 kDa (Fig 3C, red asterisk). Migration of Rpi-amr3 shifted towards ~480 kDa (Fig 3C, blue asterisk), but we did not observe co-migration of Rpi-amr3 with NRC2 oligomer (Fig 3C) at any of the time points we investigated. Our results indicate that the helper NLR NRC2 forms a high-molecular weight complex upon activation, that does not contain the sensor NLR Rpi-amr3.

**Figure 3.**
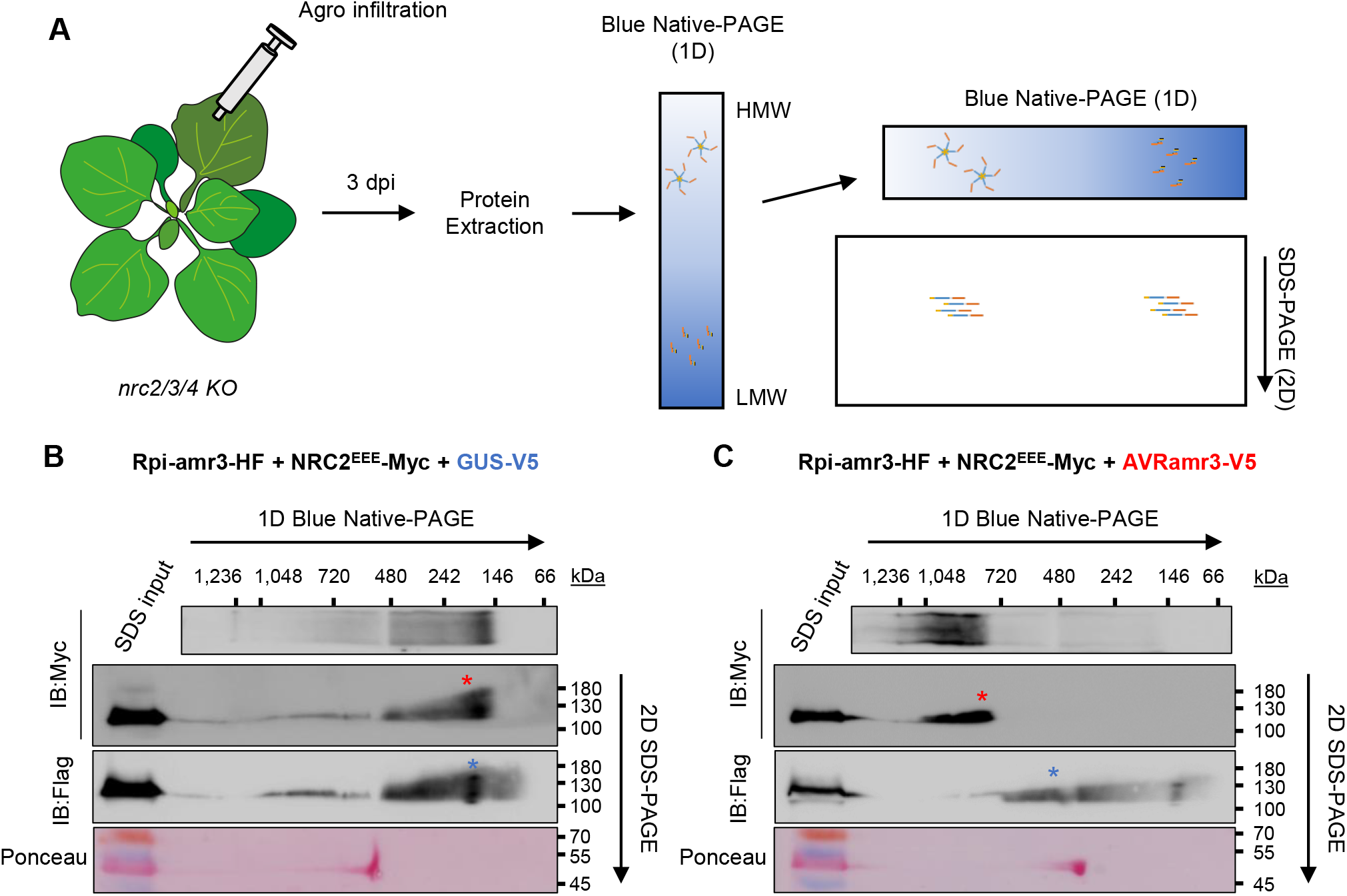
Rpi-amr3 is not present in oligomerized NRC2 protein complex. A. Experimental design for 2D-PAGE (blue native-PAGE/SDS-PAGE). Agro-infiltrated *nrc2/3/4 KO N. benthamiana* plants were collected at 3dpi for protein extraction. Protein extracts were loaded on blue native-PAGE (1D) to separate high molecular weight (HMW) protein complexes (hypothesized as a pentamer) from low molecular weight (LMW) protein complexes. Subsequently, blue native-PAGE gels were loaded on SDS-PAGE (2D) for separation of protein complexes into individual proteins. B. NRC2^EEE^ and Rpi-amr3 migrate as monomers in the absence of effector. *N. benthamiana nrc2/3/4 KO* plants were transiently infiltrated with Rpi-amr3-HF, NRC2^EEE^-Myc and GUS-V5 followed by 2D-PAGE. Molecular weight markers for blue native-PAGE are labelled on top, and SDS-PAGE markers are labelled on the right. Ponceau S staining of rubisco large subunit serves as control. NRC2^EEE^-Myc protein complex between 146~242 kDa is indicated (*). Rpi-amr3-HF protein complex between 146~242 kDa is also indicated (*). C. *N. benthamiana nrc2/3/4 KO* plants were transiently infiltrated with Rpi-amr3-HF, NRC2^EEE^-Myc and AVRamr3-V5 followed by 2D-PAGE. Molecular weight markers for blue native-PAGE are labelled on top, and SDS-PAGE markers are labelled on the right. NRC2^EEE^-Myc protein complex >720 kDa is indicated (*). Rpi-amr3-HF protein complex ~480 kDa is also indicated (*). Data information: Ponceau S staining of rubisco large subunit serves as control. Experiments were repeated at least 3 times with similar results. Protein lysates boiled in SDS were loaded on the same gel as control for size (SDS input).

### The NB-ARC domain of Rpi-amr3 is required for NRC2 oligomerization upon AVRamr3-dependent activation of Rpi-amr3

The NB-ARC (Nucleotide-binding domain shared by APAF1, R protein and CED-4) domain of NLRs is known to bind the β-phosphate group of ATP and is required for function (Tameling *et al*, 2002; Wang *et al*., 2019a; Wang *et al*, 2019b). In ZAR1, the replacement of ADP with ATP in this NB-ARC domain is crucial for oligomerization (Wang *et al*., 2019a; Wang *et al*., 2019b). We tested whether mutation of the ATP-binding activity of Rpi-amr3 impairs binding with its cognate effector AVRamr3 or with NRC2 oligomerization. The K182 residue of Rpi-amr3 lies within the P-loop and mutating this residue to alanine leads to loss of HR in the presence of AVRamr3 (Fig 4A). When Rpi-amr3-HF immunoprecipitated samples were used to perform blue native-PAGE, Rpi-amr3^K182A^-HF showed two major protein complexes, indistinguishable from wild-type Rpi-amr3 (Fig 4B). Co-immunoprecipitated AVRamr3-V5 also migrated to a similar position in the gel as Rpi-amr3-HF and Rpi-amr3^K182A^-HF (Fig 4C). Therefore, interaction between Rpi-amr3 and AVRamr3 is not impaired by mutation in the P-loop of Rpi-amr3, indicating that sensor NLR-effector interaction is necessary but insufficient for defense activation.

**Figure 4.**
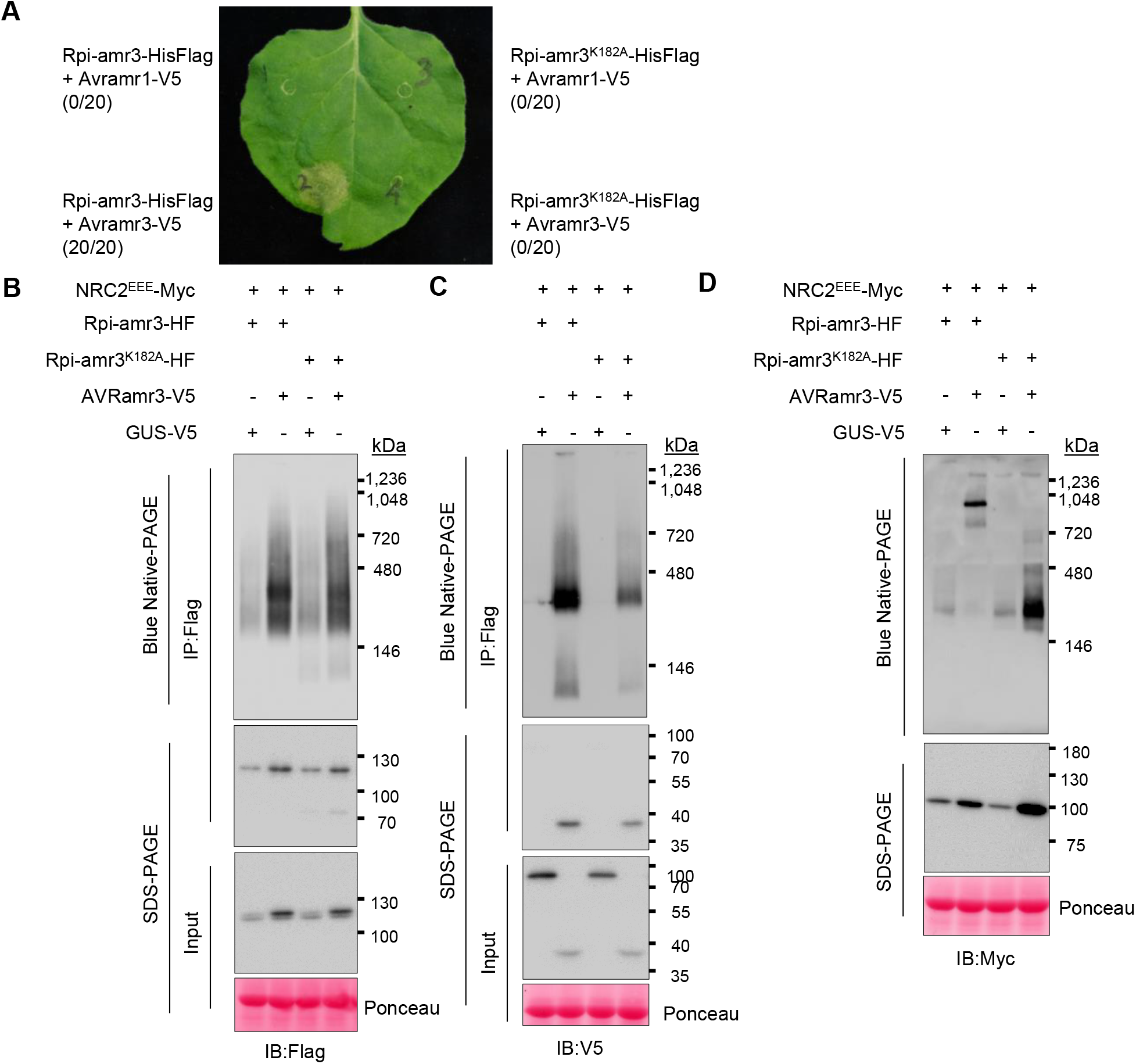
P-loop of Rpi-amr3 is required for NRC2^EEE^ oligomerization. A. P-loop of Rpi-amr3 is required for AVRamr3-dependent HR in *N. benthamiana*. Representative leaf phenotype of HR (hypersensitive response) in wild-type *N. benthamiana*. At least 20 leaves were tested, and rates of HR appearance are indicated in parentheses. B. P-loop of Rpi-amr3 is dispensable for association with AVRamr3. Protein extracts from *N. benthamiana nrc2/3/4 KO* plants were immunoprecipitated with anti-Flag antibody and blue native-PAGE was performed. Membranes were immunoblotted with anti-Flag. C. AVRamr3 associates with both wild-type Rpi-amr3 and P-loop mutant Rpi-amr3^K182A^. Protein extracts from *N. benthamiana nrc2/3/4 KO* plants were immunoprecipitated with anti-Flag antibody and blue native-PAGE was performed. Membranes were immunoblotted with anti-V5. E. NRC2^EEE^-Myc requires functional P-loop of Rpi-amr3 for oligomerization. Protein lysates from *N. benthamiana nrc2/3/4 KO* plants were used to perform blue native-PAGE. Membranes were immunoblotted with anti-Myc. Data information: SDS-boiled samples of input protein extract and IP eluates were loaded onto SDS-PAGE as control. Ponceau staining serves as loading control. Molecular weight markers are shown on the right. Experiments were done at least three times with similar results.

Next, we tested for NRC2 oligomerization upon co-expression of Rpi-amr3^K182A^-HF and AVRamr3-V5. Co-expression of AVRamr3-V5 with Rpi-amr3-HF, but not with Rpi-amr3^K182A^-HF, induces NRC2^EEE^-Myc oligomerization (Fig 4D). Therefore, while sensor NLR-effector interaction is insufficient for NRC2 activation, the ATP-binding motif of the sensor NLR is required for activating NRC2. We also observed low-abundance NRC2 forms that migrate at intermediate sizes when co-expressed with Rpi-amr3^K182A^-HF, smaller than those seen upon defense activation by a functional sensor NLR (Fig 4D). Conceivably, these might represent intermediate states of NRC2 activation.

### Multiple AVRamr3 alleles recognized by Rpi-amr3 trigger NRC2 oligomerization

AVRamr3 is a conserved RXLR effector found in multiple *Phytophthora* species. Rpi-amr3 can recognize multiple homologs of AVRamr3 from different *Phytophthora* species, including AVRamr3 from *P. parasitica* (Lin *et al*., 2021) (Fig 5A). In contrast, AVRamr3 from *P. capsici* is not recognized by Rpi-amr3 (Fig 5A, Fig EV3A) (Lin *et al*., 2021).

**Figure 5.**
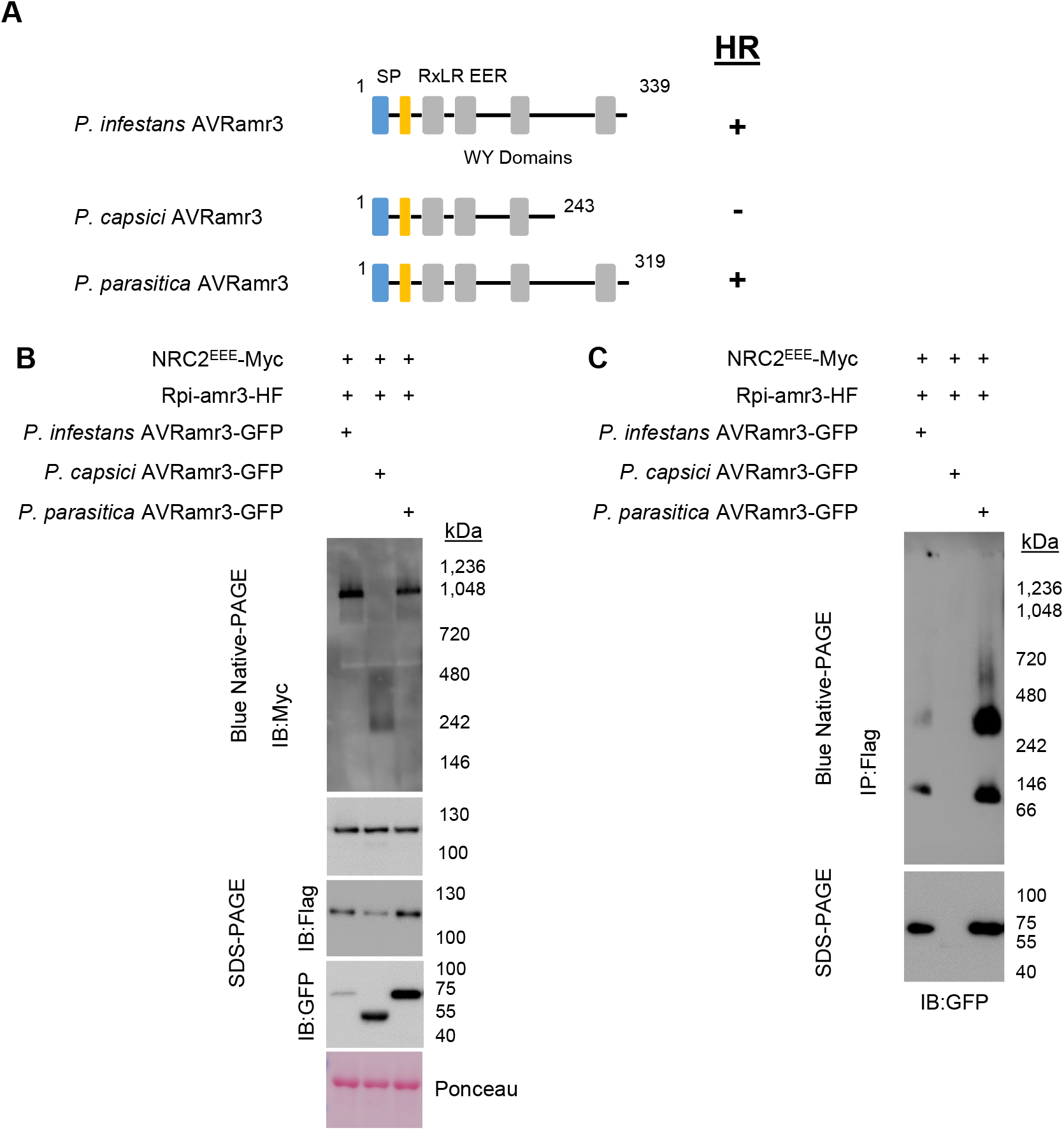
Recognized AVRamr3 alleles from different *Phytophthora* species can trigger oligomerization of NRC2^EEE^. A. Cartoon depicting different alleles of AVRamr3 from different *Phytophthora* species, *Phytophthora infestans, P. capsici*, and *P. parasitica*. Recognition of the corresponding alleles by Rpi-amr3, and thus occurrence of HR is indicated as + (recognition) or – (no recognition). B. NRC2^EEE^-Myc oligomerizes in recognition-dependent manner. Protein lysates from *N. benthamiana nrc2/3/4 KO* transiently expressing NRC2^EEE^-Myc, Rpi-amr3-HF and AVRamr3 alleles from *P. infestans, P. capsici*, and *P. parasitica* were loaded on blue native-PAGE. C. Recognition is correlated with interaction and protein complex formation of AVRamr3 with Rpi-amr3. Protein extracts from Fig 5B were immunoprecipitated with anti-Flag antibody, separated on blue native-PAGE, and AVRamr3-GFP proteins of *P. infestans, P. capsici*, and *P. parasitica* were visualized. Data information: SDS-boiled input and IP eluates were loaded onto SDS-PAGE as control. Ponceau staining serves as loading control. Molecular markers are indicated on the right. Experiments were done at least three times with similar results.

Here, to study whether other recognized AVRamr3 homolog can also activate the NRC2 resistosome, we tested for NRC2 oligomerization upon Rpi-amr3 activation with AVRamr3 alleles from different *Phytophthora* species. NRC2^EEE^-Myc oligomerizes in the presence of Rpi-amr3-HF with *P. infestans* and *P. parasitica* AVRamr3-GFP (Fig 5B) but not with *P. capsici* AVRamr3-GFP. The non-recognized truncations of AVRamr3 allele AVRamr3-T9-V5 (Fig EV5A) also did not induce NRC2 oligomerization (Fig EV5B). This shows that NRC2 oligomerization always results upon recognition of AVRamr3 alleles by Rpi-amr3.

Previously, we showed that the recognized alleles of AVRamr3-GFP from *P. parasitica* could be co-immunoprecipitated by Rpi-amr3-HF, whereas AVRamr3-GFP from *P. capsici* could not be co-immunoprecipitated (Lin *et al*., 2021). To test whether recognized alleles of AVRamr3 can form a complex with Rpi-amr3, Rpi-amr3-HF was immunoprecipitated from samples co-expressing AVRamr3-GFP from *P. parasitica*, and *P. capsici*. When these immunoprecipitates were analyzed by blue native-PAGE, *P. infestans* AVRamr3-GFP and *P. parasitica* AVRamr3-GFP migrated similarly, at both ~66 kDa and ~300 kDa (Fig 5C, Fig EV5C). However, no signals were detected for *P. capsici* AVRamr3 on blue native-PAGE, as Rpi-amr3-HF could not co-immunoprecipitate the *P. capsici* allele of AVRamr3 (Fig 5C, Fig EV5C). Rpi-amr3-HF co-expressed with non-recognized alleles of AVRamr3 also migrated as single protein species on blue native-PAGE (Fig EV5D), further confirming that non-recognized homolog or truncation of AVRamr3 do not interact with Rpi-amr3.

## Discussion

Here, we report that effector-dependent sensor NLR activation leads to oligomerization of the NRC2 helper NLR. NRC2 oligomerization is dependent on a functional P-loop in the sensor NLR, and on the cognate recognized effector. However, at the time points we analyzed, the sensor NLR does not itself oligomerize upon interaction with the recognized effector. This indicates an important difference in mode of activation between the sensor NLR and the helper NLR.

We used here the transient expression of the *S. americanum* sensor NLRs Rpi-amr1 and Rpi-amr3 with their cognate effectors AVRamr1 and AVRamr3 from *P. infestans*. The *N. benthamiana* NRC proteins can support HR upon AVRamr3 recognition by Rpi-amr3 and AVRamr1 recognition by Rpi-amr1 (Lin *et al*., 2021; Witek *et al*., 2021). To prevent cell death upon effector-dependent sensor NLR activation, we used *N. benthamiana* NRC2 mutated within its N-terminal MADA motif. NRC4^L9E^ mutated at the MADA motif activated either in an effector-dependent manner or by autoactive mutation forms punctate structures in the cell membranes (Duggan *et al*., 2021). This implies that the N-terminal MADA motif mutation of L9E/L13E/L17E of NRC2 is likely to only affect any possible channel activity and prevent cell death but should not compromise oligomerization. Contreras *et al*. (2022) show that activation of NRC2^EEE^-Myc by Rx results in enhanced association with the membrane fraction.

We investigated changes in the properties of sensor and helper NLRs using blue native-PAGE. This method uses the property of Coomassie G-250 to bind to the hydrophobic surfaces and basic amino acid residues of protein complexes without dissociating the proteins (Wittig *et al*, 2006). Due to this binding, the effect of isoelectric point of protein complexes on migration within the polyacrylamide gel becomes negligible (Wittig & Schagger, 2008). This enables the resolution of different protein complexes based on their size. However, the binding capacity of Coomassie G-250 may vary depending on protein properties, and therefore estimation of the mass of the bands observed in the blue native-PAGE is not completely reliable (Wittig *et al*., 2010). Nevertheless, blue native-PAGE is a useful method that enables detection of significant changes in molecular weight of protein complexes, as we showed for NRC2 and for the Rpi-amr3/AVRamr3 complex. Furthermore, using blue native-PAGE, we were able to distinguish effector-independent Rpi-amr1 and Rpi-amr3 self-association from the effector-bound Rpi-amr1 or Rpi-amr3 protein complexes (Fig 1). This highlights the utility of blue native-PAGE in distinguishing heterogeneous protein complexes *in vivo*.

NRC helper NLRs contain the conserved MADA motif also found in ZAR1 (Adachi *et al*., 2019). Many sensor NLRs in the NRC-dependent superclade lack the MADA motif including Rpi-amr1 and Rpi-amr3 (Witek *et al*., 2016; Witek *et al*., 2021). The MADA motif of ZAR1, which also overlaps with the α-helix that protrudes upon effector-dependent conformational change, is required for the funnel formation that leads to membrane localization and induction of cell death (Wang *et al*., 2019a). Therefore, absence of the MADA motif in sensor NLRs suggests that sensor NLRs do not participate in resistosome formation and in potential channel formation in the membrane.

Rpi-amr1 and Rpi-amr3 belong to the NRC-dependent superclade and both can use NRC2 to execute cell death and resistance (Witek *et al*, 2021; Lin *et al*, 2021). We found that both Rpi-amr1 and Rpi-amr3 interact with their cognate effector independent of NRC2, and this interaction with cognate effector induces NRC2 oligomerization. The NRC2^EEE^-Myc oligomerization patterns resulting from Rpi-amr1 and Rpi-amr3 activation were indistinguishable (Fig 2C, D). In a companion paper (Contreras *et al*, 2022), the NRC-dependent CC-NLRs Bs2 and Rx were also shown to activate NRC2 oligomerization into slower migrating forms indistinguishable from what we report here. This indicates that NRC2 oligomerization to ~900 kDa is a universal mode of action for NRC-dependent Solanaceae sensor NLRs, and further supports the conclusion that sensor NLRs are not included in the NRC2 resistosome. Our analysis with 2D-PAGE provides further evidence that Rpi-amr3 is not incorporated in the NRC2 resistosome (Fig 3C).

The NB-ARC domain is required for NLR function (Tameling *et al*., 2002). The ATP-binding P-loop motif is known to be required for activation of Rx (Bendahmane *et al*, 2002), another NRC-dependent sensor NLR. We found that similar to Rx, the P-loop motif of Rpi-amr3 is required for cell death. Oligomerization of NRC2^EEE^ in the presence of AVRamr3 and Rpi-amr3 was lost when we tested the P-loop mutant of Rpi-amr3 (Rpi-amr3^K182A^) (Fig 4D). However, we also found that Rpi-amr3^K182A^-HF retains its interaction with AVRamr3 (Fig 4B, C). Therefore, a mutation in the P-loop region does not alter the sensor NLR’s capacity to interact with its cognate effector, but instead affects its downstream signaling that provokes oligomerization of helper NLRs. Recent structural analysis of CC-NLRs shed further light on the role of the NB-ARC domain in its oligomerization. The release of ADP in exchange for ATP is required for the NB-ARC domain-mediated packing of both ZAR1 and Sr35 molecules into resistosomes (Förderer *et al*., 2022; Wang *et al*., 2019a).

In mammalian cells, NLRC4 is activated by NAIP2-mediated recognition of bacterial flagellin, or NAIP5/6-mediated recognition of bacterial PrgJ (Zhao *et al*, 2011). The NAIP is incorporated into the inflammasome and co-migrates with NLRC4 in non-denaturing PAGE (Kofoed & Vance, 2011). Ligand-bound NAIP undergoes a conformational change that leads to interaction and subsequent intermolecular autoactivation of NLRC4 (Hu *et al*, 2015; Zhang *et al*, 2015). This may explain the near-complete conversion of NLRC4 molecules to inflammasome (Kofoed & Vance, 2011). We also observed near complete conversion of the majority of NRC2 molecules into oligomers (Fig 2C, D). This may indicate that the conformational change in NRC2 upon Rpi-amr3 activation is similar to NLRC4 oligomerization. The conformational change of NRC2 induced by Rpi-amr3/AVRamr3 protein complex may trigger self-propagation of interaction and formation of a complete NRC2 resistosome. On the other hand, in contrast to NAIP/NLRC4 inflammasome formation, our data, and the data of Contreras et al (companion paper), indicate that the activated sensor NLR is not stably incorporated into the activated NRC2 complex (Fig 2, Fig 3).

To explain these observations, we propose a simple model, in which a transient interaction of activated Rpi-amr3 or Rpi-amr1 with NRC2 converts inactive NRC2 into activated NRC2 that can activate additional NRC2 protomers, enabling NRC2 oligomerization. This model is depicted in cartoon form in Figure 6. In the pre-activated state, the sensor NLRs, such as Rpi-amr3 or Rpi-amr1, reside in the cells mainly as monomers. Conceivably, additional proteins, such as chaperones, could be bound to these monomeric NLRs. Upon secretion of effectors from various *Phytophthora* species into the plant cell, some effectors (AVRamr3 in Fig 6) interact with their recognizing sensor NLR (Rpi-amr3 in Fig 6). The formation of a stable complex between effector and NLR induces activation and/or conformational change of the sensor NLRs. Then, the activated sensor NLR interacts transiently with the helper NRC2, triggering a conformational change, perhaps in an analogous manner to that by which an activated NAIP triggers a conformational change in NLRC4, but without stably incorporating the sensor NLR. The activated NRC can interact with, activate, and oligomerize with additional NRC molecules, without the need to retain the interaction with the sensor NLR. Therefore, the transiently interacting complex of sensor and helper NLR might be expected to be very low in abundance and very short-lived. The ATP-binding P-loop motif of the sensor NLR is required for activation of NRC2, conceivably due to the conformational change of the NB-ARC domain in the sensor NLR. By analogy with ZAR1(Wang *et al*., 2019a), the activated NRC2 may form a pentamer. We were not able to observe interaction between sensor and helper NLRs at the time points we tested. This is likely due to low abundance and short half-life of these protein complexes. Moreover, we cannot exclude the possibility that there may be additional proteins or molecules that mediate signaling between sensor NLRs Rpi-amr3 (and Rpi-amr1) with helper NLR NRC2, and this possibility will be investigated in future work.

**Figure 6.**
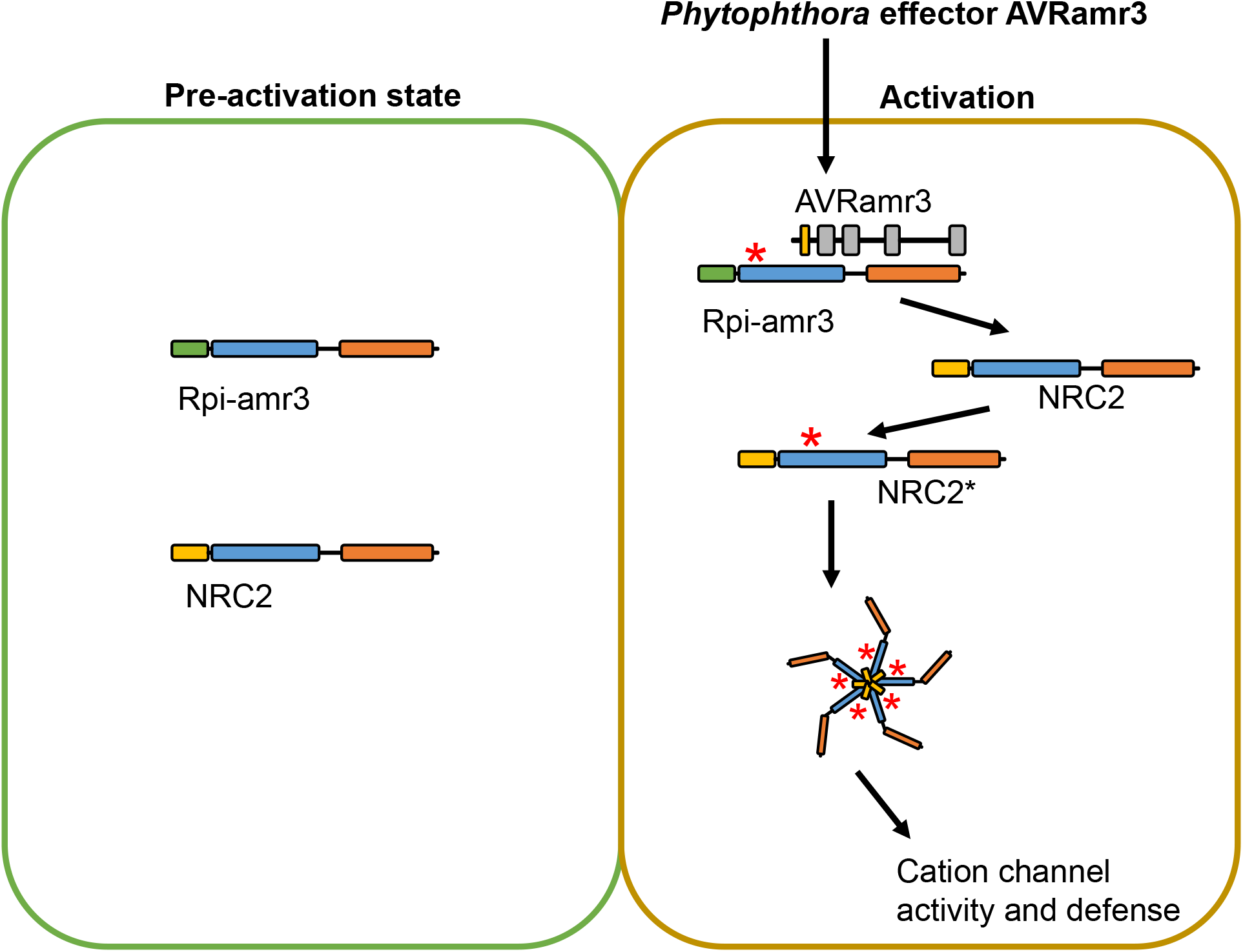
Model for activation of helper NLR NRC2 upon recognition of AVRamr3 by Rpi-amr3. Interaction with AVRamr3 converts Rpi-amr3 into an activated form (indicated by an *) that can interact with and activate NRC2 into an activated form that triggers activation of additional NRC2 protomers, enabling assembly of an NRC2 resistosome.

Previously, we reported that Rpi-amr3 can recognize AVRamr3 homologs from 9 different *Phytophthora* species, including *P. infestans*, *P. parasitica*, *P. cactorum*, *P. palmivora*, *P. megakarya*, *P. litchi*, *P. sojae*, *P. lateralis* and *P. pluvialis*. Rpi-amr3 associates only with the HR-triggering AVRamr3 homologs. We further showed that *Rpi-amr3* confers resistance against *P. infestans, P. parasitica* and *P. palmivora* in transgenic *N. benthamiana* lines (Lin *et al*, 2021). In this study, we revealed the mechanism of Rpi-amr3 activation, and found that the recognized AVRamr3 homologs from *P. infestans*, and *P. parasitica* (but not the non-recognized AVRamr3 homolog from *P. capsici)* can provoke the appearance of a ~900 kDa NRC2 resistosome through Rpi-amr3 (Fig 5B).

Unlike genes encoding surface immune receptors, e.g. *EFR*, which retain function when transferred between plant families (Lacombe *et al*, 2010), a limiting factor of NLR gene-based resistance engineering is restricted taxonomic functionality (RTF) (Tai *et al*., 1999). In many cases, *NLR*-encoding genes can only confer resistance in closely related plant species. For example, the bacterial spot disease resistance gene *Bs2* from pepper is not functional in Arabidopsis (Tai *et al*, 1999). The identification of the NRC network (Wu *et al*., 2017) and our finding on NRC activation might help to break RTF and enable breeders to deploy *NLR* genes across different plant families. Many *Phytophthora* species that carry recognized AVRamr3 infect plant species that lack *NRC* genes, such as *P. palmivora* infecting cacao, *P. cactorum* infecting strawberry and *P. sojae* infecting soybean. Co-delivery of *Rpi-amr3* and *NRC* genes into these plants might help to protect them against these *Phytophthora* pathogens.

## Materials and Methods

### Plant materials and growth conditions

The wild-type *Nicotiana benthamiana* and *NRC2*, *NRC3* and *NRC4* knockout *N. benthamiana* line *nrc2/3/4.210.4.3* were used in this study (Wu *et al*., 2020). The plants were grown in a controlled environment room (CER), with 16 hours photoperiod, at 22 °C and 45-65% humidity.

### Constructs

To clone the genes with different tags, all the ORFs (open reading frames) without stop codon were cloned into a golden gate compatible level 0 vector (pICSL01005). Then these were fused with different C-terminus tags and shuffled into a binary vector pICSL86977OD (with 35S promoter and Ocs Terminator). The C-terminus tags used in this study are C-HisFlag (PICSL50001), C-V5 (PICSL50012), C-Myc (PICSL50010), C-HA (PICSL50009), C-GFP (PICSL50008), C-3xFlag (PICSL50007), NLUC-Flag (pICSL50047) and CLUC-Flag (pICSL50048). All the constructs used in this study are listed in Table S1. The NRC^EEE^-Myc construct was cloned into pJK268c (Kourelis *et al*, 2020).

### Agrobacterium infiltration

The binary constructs were transformed into *Agrobacterium* strain GV3101-pMP90 and stored in a −80 °C freezer with 20% glycerol. Two days before the *Agrobacterium* infiltration, the constructs were streaked out on solid L medium plate (with kanamycin and rifampicin) and grown in a 28 °C incubator. For the *Agrobacterium* infiltration, 1 mM acetosyringone were added into the infiltration buffer (MgCl_2_-MES, 10 mM MgCl_2_ and 10 mM MES, pH 5.6), then the *Agrobacterium* were re-suspended into infiltration buffer, the OD_600_ was adjusted to 0.5 and the infiltration was performed 1 h later. For GUS-V5, we used OD600 =0.1; When AVRamr1-V5 and AVRamr3-V5 were used in a same experiment, we reduced the AVRamr3 construct to OD600=0.2. For the co-expression experiments, the *Agrobacterium* suspension were equally mixed before infiltration.

### HR assay

For the HR assay, four-weeks old *N. benthamiana* were used, the constructs in *Agrobacterium* were infiltrated or co-infiltrated into the abaxial surface of *N. benthamiana* leaves. The HR phenotype were scored and the photos were taken 3-4 days post *Agrobacterium* infiltration (dpi).

### Split Luciferase assay

The split-luciferase assay was described previously (Lin *et al*., 2021). In brief, *p35S::Rpi-amr1-*Cluc::OcsT and p35S::*Avramr1*-Nluc::OcsT constructs were made and transformed into *Agrobacterium* strain GV3101-pMP90. p35S::*Rpi-amr3*-Cluc::OcsT and *p35S::Avramr3*-Nluc::OcsT constructs were used as controls. The constructs were expressed or co-expressed in *nrc2/3/4* knockout *N. benthamiana* plants, OD_600_=0.5. The leaves were infiltrated with 0.4 mM luciferin on 100mM sodium citrate buffer (pH 5.6) at 3dpi, then the leaves were picked for imaging with NightOWL II LB 983 in Vivo Imaging System. Two leaves were used for each test and three independent experiments were performed with same results.

### Structure prediction using AlphaFold

Protein structure of AVRamr1 was predicted using AlphaFold and ColabFold (Jumper *et al*, 2021; Mirdita *et al*, 2022), via https://github.com/deepmind/alphafold/. The structure was visualized with ChimeraX (Pettersen *et al*, 2021), developed by the Resource for Biocomputing, Visualization, and Informatics at the University of California, San Francisco, with support from National Institutes of Health R01-GM129325 and the Office of Cyber Infrastructure and Computational Biology, National Institute of Allergy and Infectious Diseases.

### Protein extraction

Agrobacterium-infiltrated leaves were sampled with cork borer at 3 dpi. Ten disks were collected for each sample and put into 2-ml eppendorf tubes with 2 tungsten beads, which were immediately frozen in liquid nitrogen. Frozen samples were then ground in Geno/Grinder^®^ (SPEX SamplePrep) at 1,200 rpm for 2 min. Protein extraction buffer (Tris-Cl pH 7.5 50 mM, NaCl 50 mM, Glycerol 10%, MgCl_2_ 5 mM, 10 mM DTT, 0.2% NP-40, protease inhibitor cocktail) was added in equal volume across samples to ensure equal protein concentration. Centrifugation was performed at 13,000 rpm for 15 min and 5 min subsequently to remove cell debris. For subsequent blue native-PAGE analysis, sample aliquots were immediately frozen in liquid nitrogen.

### Immunoprecipitation and elution

Flag-M2 beads (Sigma, A2220) were added to the protein extract after centrifugation and incubated at 4°C for 2 h. After incubation, washing step with protein extraction buffer was performed 5 times (2 times with extraction buffer containing NP-40 0.4%, then 3 times with extraction buffer containing NP-40 0.2%) to ensure removal of non-specific binding proteins. For elution of proteins, 3xFlag peptide (Sigma, F4799) were added to beads at concentration of 0.2 mg/ml and incubated for 1 h. Sample aliquots were made for blue native-PAGE analysis and were immediately frozen in liquid nitrogen.

### SDS-PAGE and immunoblot

Protein samples were incubated at 70°C for 10 min after adding 3x SDS sample buffer (stock concentration 30% glycerol, 3% SDS, 93.75 mM Tris-Cl pH 6.8, 0.06% bromophenol blue). These samples were loaded on SDS-PAGE gels (8% or 12%) and run at 90V. After dye front reached the end, these gels were transferred with TransBlot (Biorad) at conditions of 1.0 mA for 30 min onto PVDF membranes. Transferred membranes were blocked with 5% skim milk in TBST, and antibodies were added subsequently and incubated overnight at 4°C. The following antibodies were used; Flag-HRP (Sigma, A8592), Myc-HRP (Sigma, 16-213), V5-HRP (Sigma, V2260), HA-HRP (Roche, 12013819001), and GFP-HRP (Abcam, ab6663). Signals were detected using ECL substrates (Thermo Fisher, 34580). After detection, membranes were stained with Ponceau S solution (Sigma, P7170) to use as loading control. PageRuler^™^ Prestained Protein Ladder (Thermo Scientific, 26616) was used as molecular weight markers.

### Blue native-PAGE

Blue native-PAGE was performed as indicated by the manufacturer. Protein extracts and immunoprecipitation (IP) eluates were added with 4x NativePAGE^™^ Sample Buffer (Invitrogen^™^, BN2003) and NativePAGE^™^ 5% G-250 Sample Additive (Invitrogen^™^, BN2004) to a final concentration of 0.125%. Then, samples were loaded on NativePAGE^™^ Novex^®^ 3–12% Bis-Tris Gels (Invitrogen^™^, BN1001) and run at 150 V in cathode buffer containing Coomassie G-250 (by adding NativePAGE^™^ Cathode Buffer Additive to 1/200 dilution, Invitrogen^™^, BN2002 to NativePAGE^™^ Running Buffer, Invitrogen^™^, BN2001). NativeMark^™^ Unstained Protein Standard (Invitrogen^™^, LC0725) or SERVA Native Marker (SERVA, 39219.01) were loaded to predict the size of detected protein species.

### 2D-PAGE

Samples were run on blue native-PAGE and gel strips were cut for each lane. Gel strips were put in 15 ml conical tubes and incubated with 3x SDS sample buffer containing 50 mM DTT for 15 min. These gel strips are then loaded onto 8% SDS-PAGE resolution gel. Corresponding protein extracts prepared for SDS-PAGE were loaded together to serve as control. After SDS-PAGE, transfer to PVDF membranes and immunoblots were performed as described above.

## Acknowledgements

We thank members of the Jones lab and the Kamoun lab for their helpful comments and discussions. H.-K.A. thanks Adam Bentham for his support on protein structure prediction data analysis. We would like to thank Timothy Wells, from Horticultural Services at John Innes Centre, for excellent care of the plants. We also wish to thank Mark Youles in TSL Synbio for his support with Golden Gate cloning. H.-K.A. was supported by the ERC Advanced Grant “ImmunitybyPairDesign” and core funding to J.D.G.J. from the Gatsby Charitable Foundation. X.L and A.C.O.A were supported by BBSRC grant BB/P021646/1 and the Gatsby Charitable Foundation.

## Author Contributions

H.-K.A., X.L., and J.D.G.J. conceptualized, wrote, reviewed and edited the manuscript. Experiment and analysis were performed by H.-K.A., X.L. and A.C.O.A. Methodology was developed by H.-K.A., with L.D. Resources were provided by M.C., J.K., and S.K. J.D.G.J. supervised and secured funding for the project.

## Expanded View Figures

**Figure EV1.** AVRamr1 is an RxLR effector with WY domains.

A. Predicted structure of AVRamr1 indicate 3 WY domains. Protein structures were generated using AlphaFold and visualized using ChimeraX software. Confidence level of b factors are indicated in colours.

B. WY3 domain of AVRamr1 shows conserved structure as well as the “WY” residues. Conserved α-helices of WY domain and the tryptophan (220W) and tyrosine (252Y) residues that comprise the hydrophobic core of the WY domains are indicated.

C. Predicted lDDT plot for AVRamr1 structure prediction.

D. Predicted aligned error plot for AVRamr1 structure prediction.

**Figure EV2.** AVRamr1 interacts with Rpi-amr1 *in planta*.

A. Rpi-amr1 recognizes AVRamr1 and induces HR. Wild-type *N. benthamiana* plants were transiently infiltrated, and leaf samples were imaged at 5 dpi for HR. Experiment was performed at least three times with similar results.

B. Rpi-amr1 interacts with AVRamr1 *in planta*. Constructs with truncations of luciferase (Nluc or Cluc) were transiently expressed in *nrc2/3/4 KO* N. benthamiana plants and imaged at 3 dpi.

**Figure EV3.** Defense mechanisms initiated via NRC2-Myc, but not NRC2^EEE^-Myc, lead to degradation of Rpi-amr3 and AVRamr3.

A. NRC2-Myc co-expression leads to degradation of Rpi-amr3 and AVRamr3. Immunoprecipitation with anti-Flag antibody of protein extracts in *nrc2/3/4* knockout *N. benthamiana* plants. Aliquot of samples were SDS-boiled and loaded on SDS-PAGE.

B. IP Samples from Fig EV3A were loaded on blue native-PAGE. Rpi-amr3 and AVRamr3 complex is indicated (*).

Data information: Ponceau staining serves as loading control for panel A. Molecular markers are shown on the right. Similar results were observed at least three times.

**Figure EV4.** AVRamr3-V5 and NRC2^EEE^-Myc are not affected by co-expression of each other.

Protein extracts from *nrc2/3/4* knockout *N. benthamiana* plants expressing AVRamr3-V5 and/or NRC2^EEE^-Myc were loaded on blue native-PAGE. Aliquot of the protein extracts that were treated with SDS-boiling serve as control. Ponceau staining serves as loading control. Molecular markers are indicated on the right. Experiments were repeated at three times with similar results.

**Figure EV5.** Non-recognized alleles of AVRamr3 do not trigger NRC2 oligomerization and do not interact with Rpi-amr3.

A. Schematic depiction of AVRamr3 from *Phytophthora infestans*, a truncated version of AVRamr3 (AVRamr3 T9), and AVRamr3 from *P. capsici*. Recognition of the corresponding AVRamr3 proteins by Rpi-amr3, and thus occurrence of HR is indicated as + (recognition) or – (no recognition).

B. NRC2^EEE^-Myc oligomerizes in recognition-dependent manner. Protein extracts from *N. benthamiana nrc2/3/4 KO* were loaded on blue-native PAGE.

C. Recognition is correlated with interaction and protein complex formation of AVRamr3 with Rpi-amr3. Protein extracts from Fig EV3B were immunoprecipitated with anti-Flag antibody and were loaded on blue native-PAGE.

D. Recognition is correlated with interaction and protein complex formation of Rpi-amr3 with AVRamr3. Protein extracts from Fig EV3B were immunoprecipitated with anti-Flag antibody and were loaded on blue native-PAGE.

Data information: SDS-boiled input and IP eluates were loaded onto SDS-PAGE as control. Ponceau staining serves as loading control. Molecular markers are indicated on the right. Experiments were done at least three times with similar results. Ponceau loading for Fig EV5D was same as with Fig EV5C.

## Tables and their legends

**Table S1**. Constructs used in this study.

