## Supplementary figures for "Effector-dependent activation and oligomerization of NRC helper NLRs by Rpi-amr3 and Rpi-amr1"

### Figure EV1

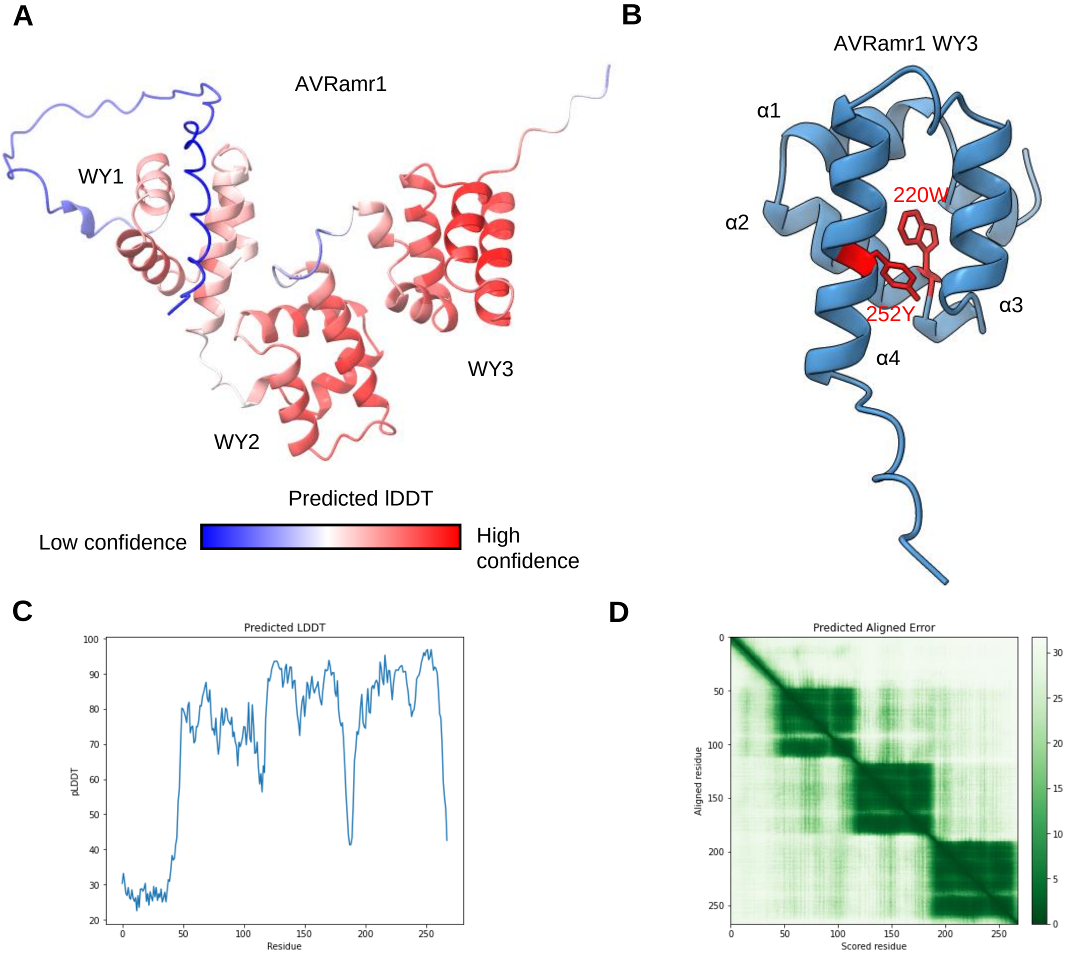


**Figure EV1.** AVRamr1 is an RxLR effector with WY domains.

A. Predicted structure of AVRamr1 indicate 3 WY domains. Protein structures were generated using AlphaFold and visualized using ChimeraX software. Confidence level of b factors are indicated in colours.

B. WY3 domain of AVRamr1 shows conserved structure as well as the “WY” residues. Conserved α-helices of WY domain and the tryptophan (220W) and tyrosine (252Y) residues that comprise the hydrophobic core of the WY domains are indicated.

C. Predicted lDDT plot for AVRamr1 structure prediction.

D. Predicted aligned error plot for AVRamr1 structure prediction.

### Figure EV2

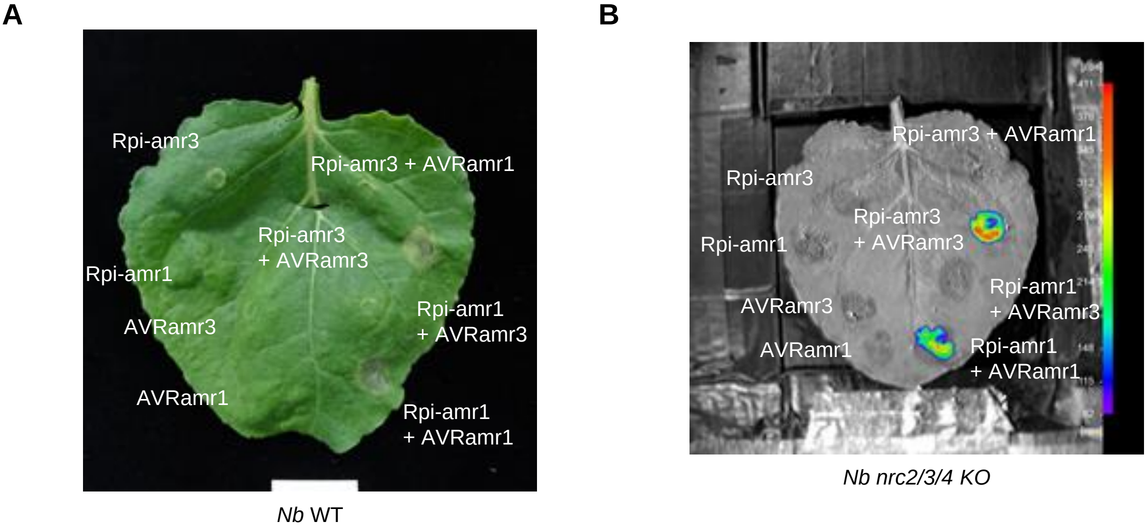


**Figure EV2.** AVRamr1 interacts with Rpi-amr1 *in planta*.

A. Rpi-amr1 recognizes AVRamr1 and induces HR. Wild-type *N. benthamiana* plants were transiently infiltrated, and leaf samples were imaged at 5 dpi for HR. Experiment was performed at least three times with similar results.

B. Rpi-amr1 interacts with AVRamr1 *in planta*. Constructs with truncations of luciferase (Nluc or Cluc) were transiently expressed in *nrc2/3/4 KO* N. benthamiana plants and imaged at 3 dpi.

### Figure EV3

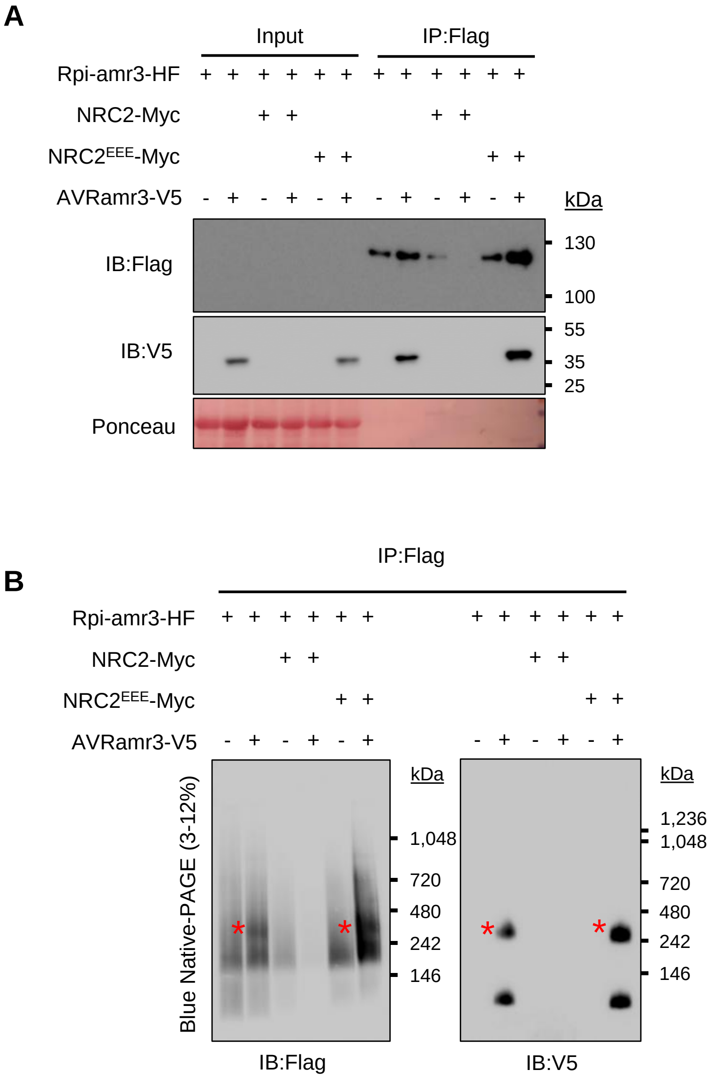


**Figure EV3.** Defense mechanisms initiated via NRC2-Myc, but not NRC2$^{EEE}$-Myc, lead to degradation of Rpi-amr3 and AVRamr3.

A. NRC2-Myc co-expression leads to degradation of Rpi-amr3 and AVRamr3. Immunoprecipitation with anti-Flag antibody of protein extracts in *nrc2/3/4* knockout *N. benthamiana* plants. Aliquot of samples were SDS-boiled and loaded on SDS-PAGE.

B. IP Samples from Fig EV3A were loaded on blue native-PAGE. Rpi-amr3 and AVRamr3 complex is indicated (*).

Data information: Ponceau staining serves as loading control for panel A. Molecular markers are shown on the right. Similar results were observed at least three times.

#### Figure EV4

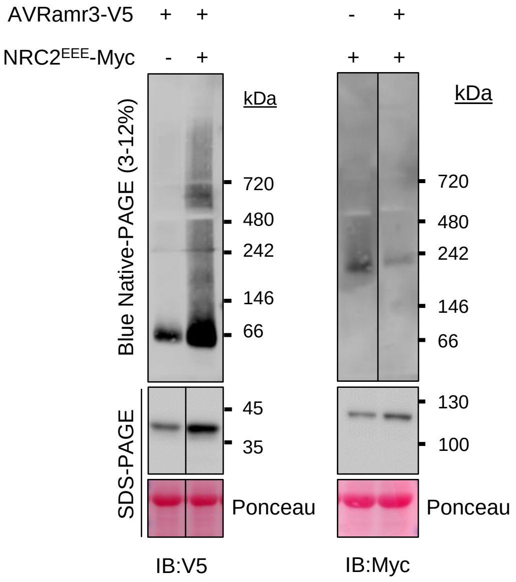


**Figure EV4.** AVRamr3-V5 and NRC2$^{EEE}$-Myc are not affected by co-expression of each other.
Protein extracts from *nrc2/3/4* knockout *N. benthamiana* plants expressing AVRamr3-V5 and/or NRC2$^{EEE}$-Myc were loaded on blue native-PAGE. Aliquot of the protein extracts that were treated with SDS-boiling serve as control. Ponceau staining serves as loading control. Molecular markers are indicated on the right. Experiments were repeated at three times with similar results.

### Figure EV5

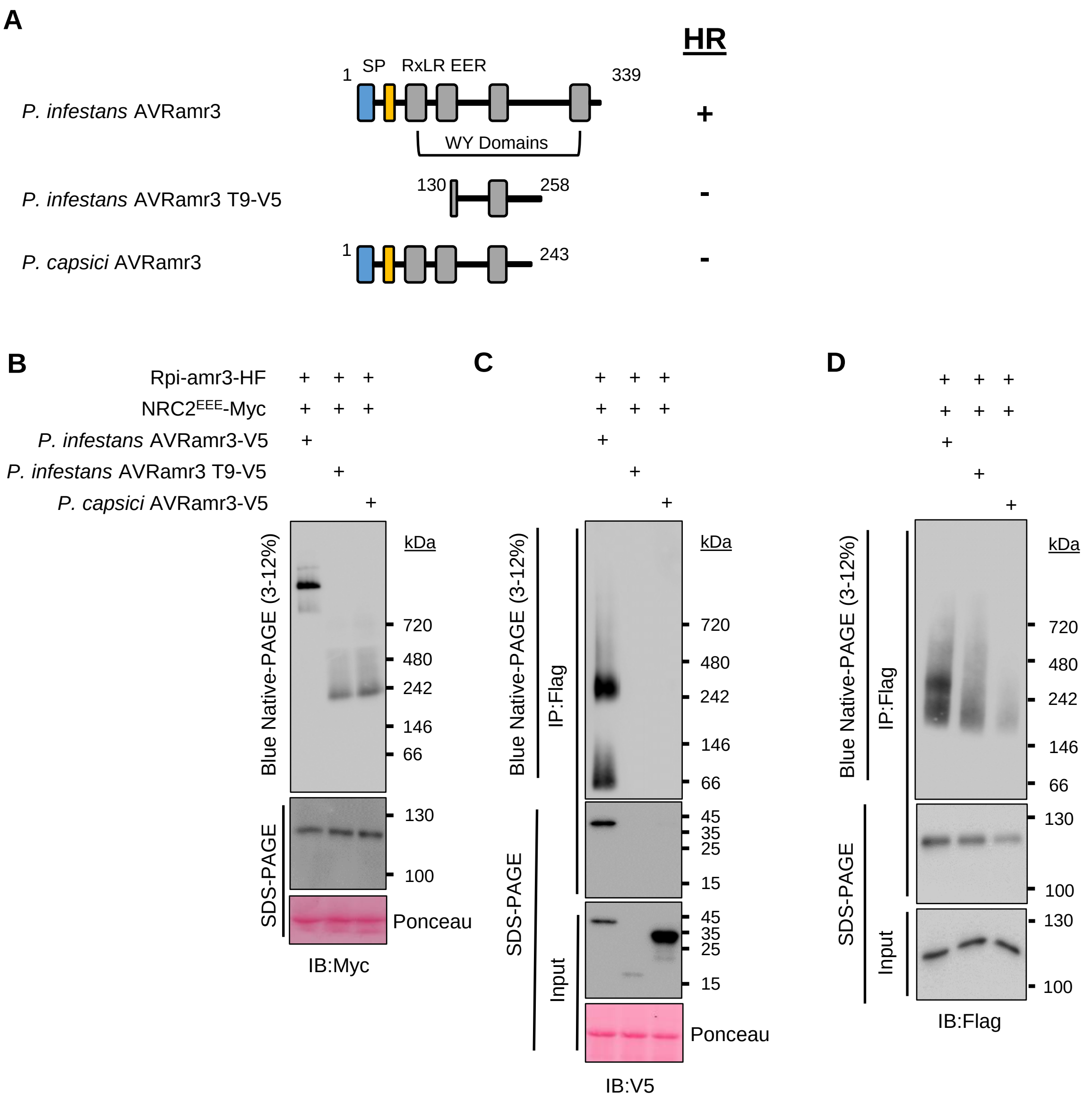


**Figure EV5.** Non-recognized alleles of AVRamr3 do not trigger NRC2 oligomerization and do not interact with Rpi-amr3.

A. Schematic depiction of AVRamr3 from *Phytophthora infestans*, a truncated version of AVRamr3 (AVRamr3 T9), and AVRamr3 from *P. capsici*. Recognition of the corresponding AVRamr3 proteins by Rpi-amr3, and thus occurrence of HR is indicated as + (recognition) or – (no recognition).

B. NRC2$^{EEE}$-Myc oligomerizes in recognition-dependent manner. Protein extracts from *N. benthamiana nrc2/3/4 KO* were loaded on blue-native PAGE.

C. Recognition is correlated with interaction and protein complex formation of AVRamr3 with Rpi-amr3. Protein extracts from Fig EV3B were immunoprecipitated with anti-Flag antibody and were loaded on blue native-PAGE.

D. Recognition is correlated with interaction and protein complex formation of Rpi-amr3 with AVRamr3. Protein extracts from Fig EV3B were immunoprecipitated with anti-Flag antibody and were loaded on blue native-PAGE.

Data information: SDS-boiled input and IP eluates were loaded onto SDS-PAGE as control. Ponceau staining serves as loading control. Molecular markers are indicated on the right. Experiments were done at least three times with similar results. Ponceau loading for Fig EV5D was same as with Fig EV5C.
